# Global monitoring of soil animal communities using a common methodology

**DOI:** 10.1101/2022.01.11.475793

**Authors:** Anton M. Potapov, Xin Sun, Maria J. I. Briones, George G. Brown, Erin K. Cameron, Jérôme Cortet, Nico Eisenhauer, Saori Fujii, Stefan Geisen, Konstantin B. Gongalsky, Carlos Guerra, I.Tanya Handa, Charlene Janion-Scheepers, Zoë Lindo, Jérôme Mathieu, Maka Murvanidze, Uffe N Nielsen, Stefan Scheu, Olaf Schmidt, Clement Schneider, Julia Seeber, Jiri Tuma, Alexei V. Tiunov, Andrey S. Zaitsev, Diana H. Wall

**Affiliations:** Johann Friedrich Blumenbach Institute of Zoology and Anthropology, University of Göttingen, Göttingen, Germany; A.N. Severtsov Institute of Ecology and Evolution, Russian Academy of Sciences, Moscow, Russia; Key Laboratory of Urban Environment and Health, Institute of Urban Environment Chinese Academy of Sciences, 361021, Xiamen, China; Departamento de Ecología y Biología Animal, Universidad de Vigo, 36310 Vigo, Spain; Embrapa Forestry, Estrada da Ribeira Km. 111, Colombo, PR, Brazil, 83411-000; Soils Department, Universidade Federal do Paraná, Rua dos Funcionários 1540, Curitiba, PR, Brazil, 80035-050; Department of Environmental Science, Saint Mary’s University, Halifax, Nova Scotia, Canada; CEFE, UMR 5175, Université Paul-Valéry Montpellier 3, Université de Montpellier, EPHE, IRD, route de Mende, 34000 Montpellier; German Centre for Integrative Biodiversity Research (iDiv), Halle-Jena-Leipzig, Puschstr. 4, 04103 Leipzig, Germany; Institute of Biology, Leipzig University, Puschstr. 4, 04103 Leipzig, Germany; Department of Forest Entomology, Forestry and Forest Products Research Institute, 1 Matsunosato, Tsukuba, Ibaraki, 305-8687, Japan; Laboratory of Nematology, Department of Plant Sciences, Wageningen University & Research, 6700 ES Wageningen, The Netherlands; Institute of Biology, Martin Luther University Halle Wittenberg, Halle (Saale), Germany; Département des sciences biologiques, Université du Québec à Montréal, CP. 8888, succursale Centre-Ville, Montréal QC, H3C 3P8, Canada; Department of Biological Sciences, University of Cape Town, Rondebosch, 7700, South Africa; Iziko Museums of South Africa, Cape Town, 8000, South Africa; University of Western Ontario, Department of Biology, London, Ontario, Canada; Institut d’Ecologie et des Sciences de l’Environnement de Paris, Sorbonne Université, CNRS, UPEC, Paris 7, INRA, IRD, F-75005 Paris, France; I. Javakhishvili Tbilisi State University. I. Chavchavadze ave. 1. 0179 Tbilisi, Georgia; Hawkesbury Institute for the Environment, Western Sydney University, Locked Bag 1797, NSW, Penrith, 2751, Australia; UCD School of Agriculture and Food Science, and UCD Earth Institute, University College Dublin, Belfield, Dublin 4, Ireland; Department of Soil Zoology, Senckenberg Society for Nature Research, Am Museum 1, 02826 Görlitz, Germany; Institute for Alpine Environment, Eurac Research, Drususallee 1, 39100 Bozen, Italy; Department of Ecology, University of Innsbruck, Technikerstrasse 25, 6020 Innsbruck, Austria; Biology Centre of the Czech Academy of Sciences, Institute of Soil Biology, Na Sádkách 7, 370 05, České Budějovice, Czech Republic; School of Global Environmental Sustainability and Department of Biology, Colorado State University, Fort Collins, CO 80523-1036, USA

**Keywords:** biogeography, ecosystem functioning, global pattern, macroecology, macrofauna, mesofauna, microfauna, soil biodiversity, soil fauna

## Abstract

Here we introduce the Soil BON Foodweb Team, a cross-continental collaborative network that aims to monitor soil animal communities and food webs using consistent methodology at a global scale. Soil animals support vital soil processes via soil structure modification, direct consumption of dead organic matter, and interactions with microbial and plant communities. Soil animal effects on ecosystem functions have been demonstrated by correlative analyses as well as in laboratory and field experiments, but these studies typically focus on selected animal groups or species at one or few sites with limited variation in environmental conditions. The lack of comprehensive harmonised large-scale soil animal community data including microfauna, mesofauna, and macrofauna, in conjunction with related soil functions, limits our understanding of biological interactions in soil communities and how these interactions affect ecosystem functioning. To provide such data, the Soil BON Foodweb Team invites researchers worldwide to use a common methodology to address six long-term goals: (1) to collect globally representative harmonised data on soil micro-, meso-, and macrofauna communities; (2) to describe key environmental drivers of soil animal communities and food webs; (3) to assess the efficiency of conservation approaches for the protection of soil animal communities; (4) to describe soil food webs and their association with soil functioning globally; (5) to establish a global research network for soil biodiversity monitoring and collaborative projects in related topics; (6) to reinforce local collaboration networks and expertise and support capacity building for soil animal research around the world. In this paper, we describe the vision of the global research network and the common sampling protocol to assess soil animal communities and advocate for the use of standard methodologies across observational and experimental soil animal studies. We will use this protocol to conduct soil animal assessments and reconstruct soil food webs on the sites included in the global soil biodiversity monitoring network, Soil BON, allowing us to assess linkages among soil biodiversity, vegetation, soil physico-chemical properties, and ecosystem functions. In the present paper, we call for researchers especially from countries and ecoregions that remain underrepresented in the majority of soil biodiversity assessments to join us. Together we will be able to provide science-based evidence to support soil biodiversity conservation and functioning of terrestrial ecosystems.

## A need for global and comprehensive soil animal ecology

Soil animals are an essential component of virtually all terrestrial ecosystems (Fierer et al., 2009; Petersen & Luxton, 1982). They support ecosystem functions by direct contribution to decomposition and nutrient cycling, and indirectly through the engineering activities, as well as changing microbial communities and plant growth (Briones, 2014; Handa et al., 2014; Hassall et al., 2006; Lavelle et al., 2006). In exclusion experiments, the presence of soil animals can enhance aboveground plant productivity by up to 70%, depending on the vegetation type (Sackett et al., 2010; Trap et al., 2016; van Groenigen et al., 2014), or facilitate litter decomposition by up to 50%, depending on climatic conditions (García-Palacios et al., 2013). These effects largely emerge from trophic and other biological interactions among key functional groups of soil animals, microbes, and plants (Bonkowski et al., 2009; Coulibaly et al., 2019; A. Potapov, 2021). Local variations in animal communities may have large effects on ecosystem processes at local, landscape, and global scales (Handa et al., 2014; Seibold et al., 2021). There were several calls to include soil animal effects in global biogeochemical (Deckmyn et al., 2020; Filser et al., 2016; Soong & Nielsen, 2016) or soil erosion models (Orgiazzi & Panagos, 2018), but the required large-scale comprehensive community data to validate these animal-based models are lacking.

The most comprehensive overview of the contribution of soil animal communities to ecosystem functioning comes from the International Biological Programme and dates back to the 1980s (Huhta, 2007; Petersen & Luxton, 1982). However, the observational data compiled at that time was not linked to soil properties and functions in a spatially-explicit way, which limited their use for biogeochemical modelling and for a broader ecosystem level understanding. Several recent studies have collected global spatially-explicit data and extrapolated global distributions of earthworms (Phillips et al., 2019) and nematodes (van den Hoogen et al., 2019), while syntheses on springtails (#GlobalCollembola) (Potapov et al. 2020; Potapov et al. 2022) and soil macrofauna (GlobalSOilMacrofauna) (Lavelle et al., *under review*) are in progress. These studies showed that the distribution of the local diversity of soil animals differs strongly from that of aboveground organisms (Cameron et al., 2019; Phillips et al., 2019) due to their contrasting responses to environmental drivers (Bardgett and van der Putten 2014). To date, global assessments of soil micro-, meso- and macrofauna have been done independently by different expert communities and using different methodologies. This limits our understanding of relationships among key functional groups of soil organisms and prevents us from global upscaling of soil communities and food webs they form. Global extrapolations of soil biodiversity were done only for a few taxonomic groups and remain poorly linked to soil functions with only 0.3% of soil ecological studies simultaneously assessing soil biodiversity and functions (Guerra et al., 2020). This important knowledge gap makes a robust quantification of soil animal contribution to biosphere functioning impossible and hampers projections of soil functioning under future global change scenarios (Guerra, Delgado-Baquerizo, et al., 2021). Furthermore, the current knowledge and especially the poorly described diversity of tropical soil communities, severely limits our understanding of human impact on soil animals and the design of appropriate conservation strategies (Eisenhauer et al., 2019; Geisen, Wall, et al., 2019; Guerra, Bardgett, et al., 2021). These knowledge gaps call for a comprehensive soil animal biodiversity assessment at a global scale using a common methodology (Eisenhauer et al., 2021; Geisen, Briones, et al., 2019; White et al., 2020), which is achievable only through a major joint effort.

### Global monitoring network

In 2018, The Soil Biodiversity Observation Network (Soil BON) was launched as a part of The Group on Earth Observations Biodiversity Observation Network (GEO BON), a United Nations initiative that aims to monitor Earth’s biodiversity (Guerra, Bardgett, et al., 2021; Scholes et al., 2008). Soil BON is a collaborative network supported by the Global Soil Biodiversity Initiative (GSBI) that focuses on soil biodiversity (Guerra, Wall, et al., 2021) and at present includes research teams across 90 countries. Local teams will take soil samples at approximately 1,000 sites across all continents except Antarctica for the first time in 2022. Additional sampling is planned every three years to establish long-term global-scale monitoring of soil biodiversity. The resulting soil samples will be shipped to a central hub (German Centre for Integrative Biodiversity Research (iDiv, Leipzig, Germany) to perform measurements of soil properties (water holding capacity, stoichiometry of nutrients, pH, root characteristics), microbiome (eDNA sequencing of prokaryotes, fungi, and protists), and functions (soil respiration, substrate-induced respiration, microbial biomass, enzymatic activity, litter decomposition, soil aggregate stability). The sampling also includes nematodes from topsoil. However, this approach is not well-suited to assess in full soil animal communities due to the large amount of materials that need to be transported (kilograms of soil per site), and the high mortality of soil animals (particularly macrofauna) in long-term stored and shipped soil samples. Hence, it is important that soil animals are collected and extracted close to the sampling site in a relatively short period of time.

Here, we introduce the Soil BON Foodweb Team (SBF Team) that focuses on the assessment of soil animal communities (Fig. 1a). We aim at producing new harmonised global data on densities and biomasses of all major taxonomic groups of soil invertebrates across micro-, meso-, and macrofauna, and thus expand the scope of the Soil BON initiative by linking soil biodiversity to ecosystem level processes through a soil food web perspective. This effort widens the core Soil BON network by involving local research communities of soil zoologists and taxonomists and applying additional sampling approaches. Instead of having a central hub, we will coordinate local researchers and facilities to form a complementary global network aiming at six main long-term goals:

**Fig. 1.**
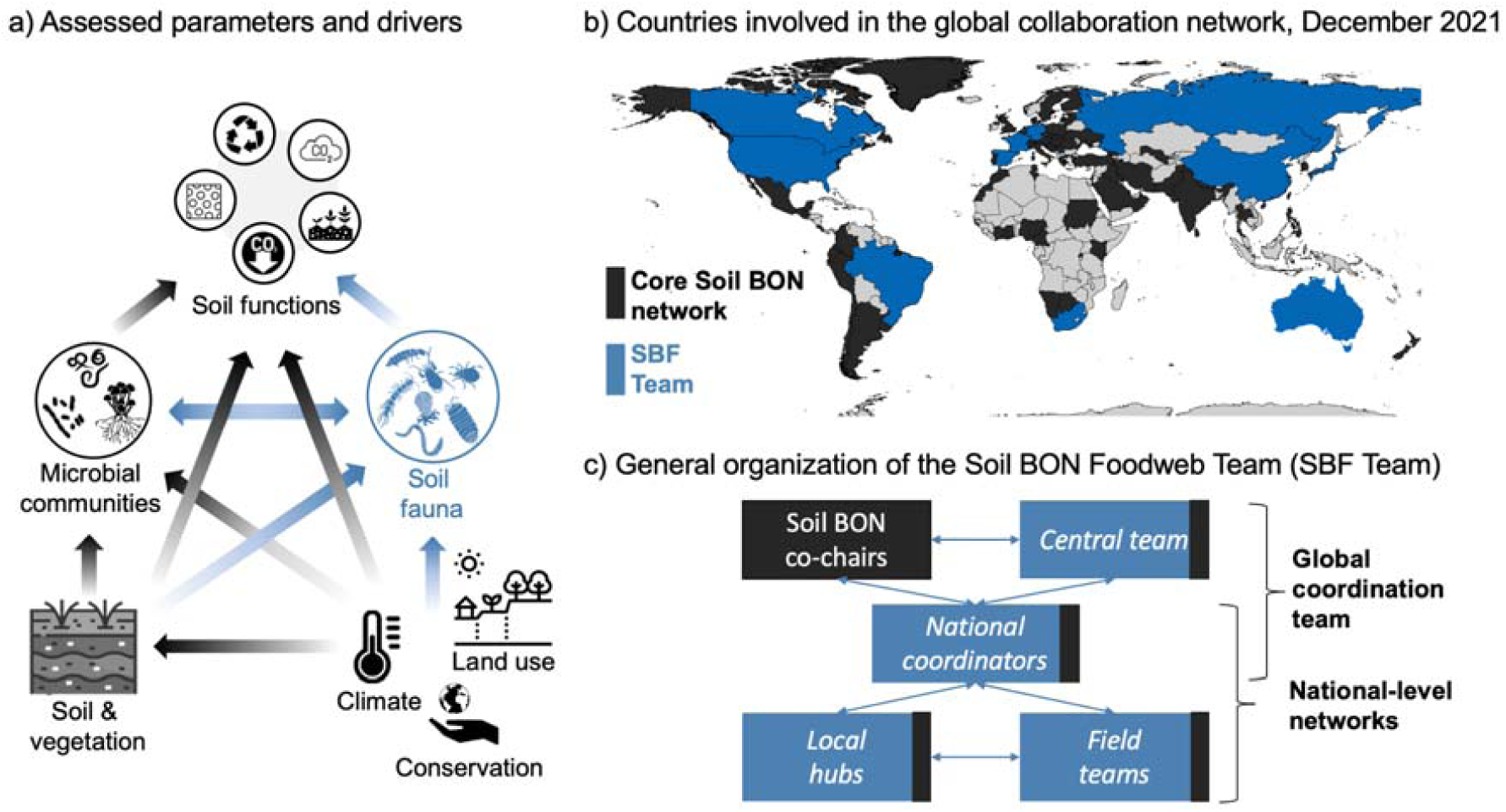
The Soil BON Foodweb Team concept. Effects of climate, land use and conservation on soil functions are mediated by soil properties, microbes, and animals; the latter are assessed by the SBF Team (a). At present, research groups from 90 countries are officially involved in Soil BON (black); the *Global coordination team* of the SBF Team (blue) covers all major continents (b). The *Global coordination team* includes the SBF *Central team*, Soil BON co-chairs, and SBF *National coordinators*. National-level networks are coordinated by *National coordinators* and may also include teams involved in the core Soil BON network (c).

1. To deliver ***open, comprehensive, and globally representative methods and harmonised datasets*** on soil micro-, meso-, and macrofauna in conjunction with soil functions.
2. To explore effects of ***climate, land use, and other environmental variables*** on soil animal communities, soil food web structure, and resistance.
3. To assess the efficiency of current ***nature conservation*** approaches in protecting soil animal communities.
4. To relate soil animal communities and food webs with ***soil functioning*** across climates, soil and land use types.
5. To establish a ***global soil fauna expert network*** for soil biodiversity monitoring and other collaborative projects.
6. To reinforce local collaboration networks and ***local expertise*** in soil zoology.

Our initial objective is to sample at least 200 out of the planned ∼1,000 Soil BON sites in 2022. Collected soil animal data will be linked to the core Soil BON network data on climate, land use, soil parameters, microorganisms, and functions via common sampling sites and similar sampling periods. Therefore, ***this is the first initiative linking quantitative soil animal data across the size spectrum to a range of soil functions worldwide***. The *Global coordination team* includes the *Central team* who is responsible for the data acquisition and storage, the co-Chairs of Soil BON, and the *National coordinators* who communicate with the local survey networks (Fig. 1c). The extended network includes *Field teams* and *Local hubs*, i.e. research teams that have expertise in soil zoology and perform soil sampling and animal extraction, respectively. All these roles are not mutually exclusive (e.g. a person can be involved in the *Global coordination team*, but also form a *Local hub* and a *Field team*). To date, research teams from 20 countries have volunteered to join the Team, partly covering also less explored regions such as Africa, South America, Asia and Russia. This paper is a call for soil animal ecologists from around the world to join the SBF Team. Priority will be given to research teams from undersampled countries and ecoregions; at present, these include mainly countries in Africa, Asia, South America, and the Middle East (Fig. 1b). To participate as a *Field team*, expertise in soil zoology, basic equipment for the field work (Supplementary protocol), and possibility to collect soil animals from several sampling sites with standardised methods is needed. To establish a *Local hub*, a laboratory equipped with either wet or dry extractors is needed. This equipment can be also built with little monetary costs and guidelines will be provided by the Team participants (Edwards, 1991; Niva et al., 2015). *National coordinators* will help create, support, and coordinate local collaboration networks of *Field teams* and *Local hubs*. The current list of *National coordinators* can be found on the web page of the SBF Team (www.soilbonfoodweb.org). All material and data contributors and national coordinators are invited to join collaborative publications, workshops, and add-on projects of the SBF Team and synthesis publications of the Soil BON consortium including their data. Below, we describe our sampling protocol, data acquisition and storage strategies.

## A common methodology

### Target variables

In the SBF Team, we aim to be ***inclusive*** for research groups and countries with limited facilities (Maestre & Eisenhauer, 2019). We also aim to be ***globally representative***, thus sampling multiple sites in different countries and environmental conditions. Finally, we want to be as ***comprehensive*** as possible and cover all key functional groups of soil micro-, meso- and macrofauna. This is a bold and challenging task that demands low-cost and labour-efficient approaches. To ensure feasibility, at the initial stage we will focus on biomass of taxonomic and functional groups, rather than on species richness. The latter implies a need for taxonomic expertise in all major soil invertebrate groups which is limiting particularly in tropical regions, where a large proportion of the individuals collected are singletons, and where many species are unknown/undescribed (Barnes et al., 2014; Demetrio et al., 2021; A. M. Potapov et al., 2020; Rossi et al., 2006). Nevertheless, building taxonomic expert capacities locally is one of the long-term goals of the SBF Team (see above). We realise that we will not be able to develop a sampling protocol that will be comprehensive enough to assess all animal groups and all research questions while keeping the workload feasible. Below, we describe the sampling approach that represents the compromise put forward by the SBF Team.

### Site selection, sampling size and time

Site selection is linked to the Soil BON core design, where soil parameters, functions, and microbial communities are assessed. If a country is included in Soil BON, sampling sites for animal assessment are chosen from the already pre-selected set. If a country is missing from the Soil BON network (Fig. 1b), the first step would be to register it to be officially enrolled (https://members.geobon.org/pages/soil-bon). The core design includes sites outside and inside areas that have a ‘protection’ status (e.g. nature conservation reserves) in each country to evaluate the effect of present conservation strategies on soil biota (Guerra, Wall, et al., 2021). The sampling of soil animals should be done at least on two sites by each *Field team* (e.g. one site inside and one outside of a protected area within the same geographical region and habitat type), but ideally cover more ecosystems and geographic areas. The final site selection is designed by *National coordinators* to identify priority ecosystems that cover wide environmental gradients (Guerra et al., 2020). At the initial stage, we target to sample at least 200 sites globally, which will be sufficient for robust general analyses (Delgado-Baquerizo et al., 2020). For all sites we collect information on the sampling date, person, location, vegetation characteristics, and history. The sampling is done at the peak vegetation biomass season according to the Soil BON approach (Guerra, Wall, et al., 2021). A 3-month window will be provided for each site and these time frames will vary across sites. At each site, soil animals are assessed at five sampling points (within a 450 m^2^ area) to account for within-site variation as the recommended minimum for large soil animals, whose spatial distribution is commonly heterogeneous (Fig. 2) (Nuria et al., 2011; Rossi et al., 2006). All samples will be processed separately, making it possible to assess local beta-diversity. Most of the field activities will be done outside sampling sites to minimise disturbance.

**Fig. 2.**
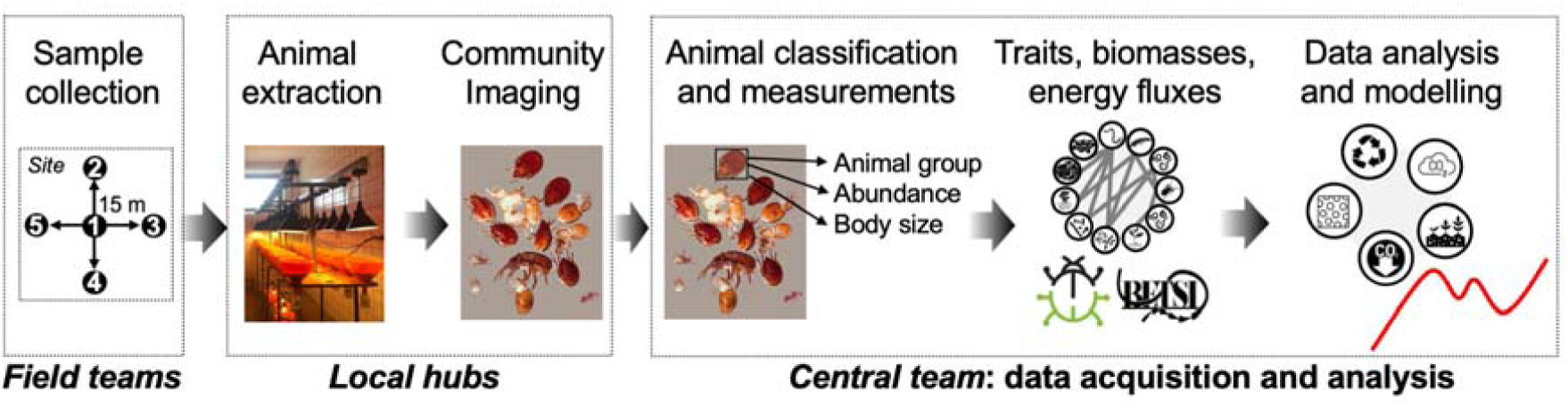
Workflow of the global soil animal assessment. Soil animals are collected by local *Field teams* at the Soil BON sites. Animals are extracted and photographed in mixed samples at *Local hubs*. Abundances and body sizes of taxonomic groups are estimated by the *Central team* using manual image annotations and a trained image analysis algorithm (Sys et al., *under review*). Biomasses and energy fluxes are estimated using allometric regressions and food-web reconstruction approaches (Ehnes et al., 2011; Jochum et al., 2021; A. Potapov, 2021; Sohlström et al., 2018) and used in statistical analyses and modelling.

### Field sampling methods

#### Selected methods

We will use a combination of sampling methods to comprehensively represent soil animal communities, including micro-, meso-, and macrofauna (Fig. 3) (Geisen, Briones, et al., 2019; White et al., 2020). Microfauna and enchytraeids will be collected using wet extraction (Baermann funnels; Niva et al, 2015; Cesarz et al., 2019) microarthropods using dry extraction (e.g. Berlese-like; (Edwards, 1991; Moreira et al., 2012), and macrofauna using hand-sorting (Anderson & Ingram, 1993; Bignell, 2009) (details are given below). We have chosen these sampling methods as the most commonly used for the corresponding animal size groups and are straightforward enough to be used even by researchers with limited experience in soil ecology. These methods provide area-based estimations of biomass and density which is necessary for data comparability and calculations of energy fluxes and functional impacts of soil animals (Jochum & Eisenhauer, 2021). For the latter reason, pitfall traps are not included in the main methods, but suggested as an auxiliary method (Supplementary protocol). We will also take pictures of topsoil profiles and sampling sites to collect environmental data (Fig. 3; Supplementary protocol).

**Fig. 3.**
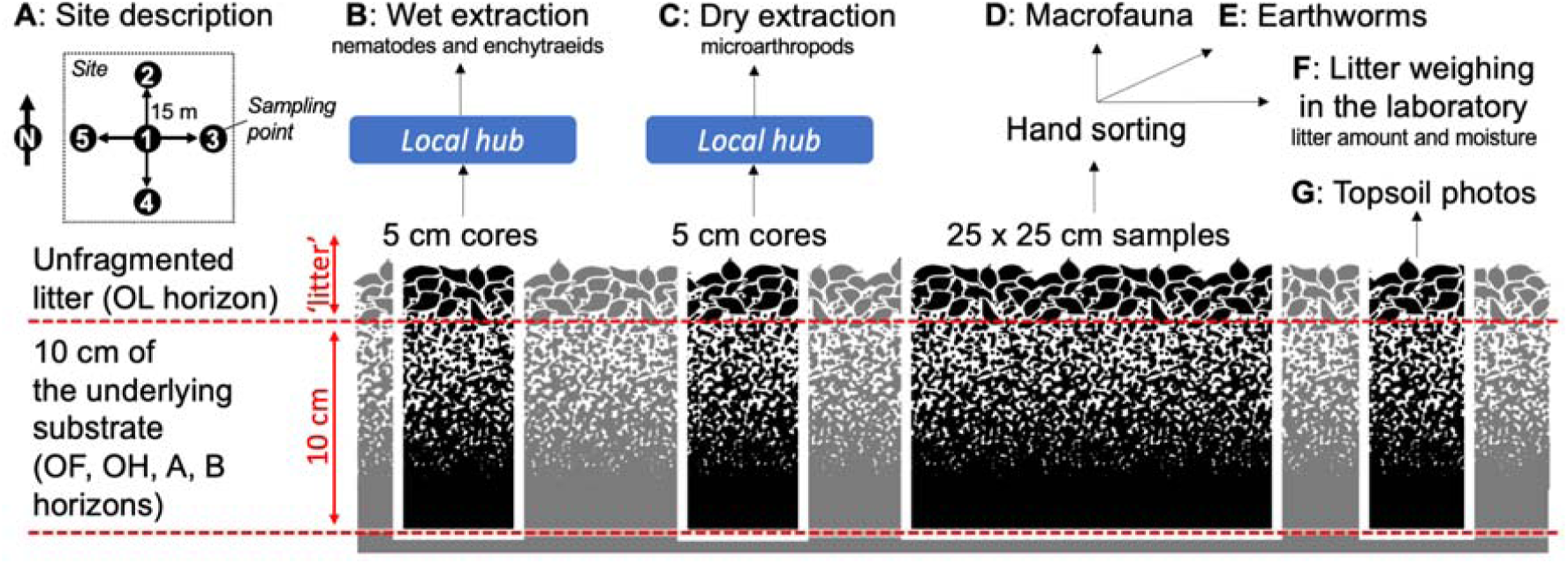
Sampling concept and animal extraction methods. A combination of wet extraction, dry extraction, and hand-sorting is used to collect soil micro-, meso-, and macrofauna. Five samples of each type are taken per *site*. Sample depth (litter and 10 cm of the underlying soil) has been chosen to ensure comparability of animal communities data with data on microorganisms, soil parameters, and functions measured in the first 10 cm of soil by the core Soil BON network. Note that we define ‘litter’ to include the OL horizon only, i.e. unfragmented leaves (Zanella, Ponge, Gobat, et al., 2018; Zanella, Ponge, Jabiol, et al., 2018). Extractions of microarthropods, nematodes and enchytraeids are done at *Local hubs*.

#### Sample areas

We follow a common 19.6 cm^2^ sample area for microarthropods, and for microfauna and enchytraeids (soil corer of 5 cm diameter; also used in the core Soil BON network sampling), and 625 cm^2^ for macrofauna (soil monoliths 25 × 25 cm). Although a larger sample area would better represent large-sized actively-moving animals and social insects, we have to limit collection efforts to make the sampling of many sites feasible and thus make the initiative more inclusive for research teams and globally representative.

#### Sample depth

We sample the entire fresh litter layer (OL horizon, unfragmented litter) together with the first 10 cm of the underlying substrate (‘soil’, here referring to OF, OH, and A horizons). Animals from litter and soil layers are collected and processed together to maximise efficiency and avoid data mismatch due to ambiguities in the definition of ‘litter’ (Fig. 4). Our assessment will focus on animals and processes in top soil and will miss deep-living soil animals (e.g., some endogeic earthworms; Lavelle, 1988), especially in soils with well-developed organic horizons (A. M. Potapov et al., 2017). However, this way would allow the application of a single standard protocol and increase the number of sites to make the analysis globally representative. The uniform sampling depth is used for all samples to make the energy flux calculations comparable across size classes. To correctly measure the sampling depth, we strongly encourage soil zoologists to become familiar with the recent HUMUSICA publications to define the diagnostic horizons (Zanella, Ponge, Gobat, et al., 2018; Zanella, Ponge, Jabiol, et al., 2018).

**Fig. 4.**
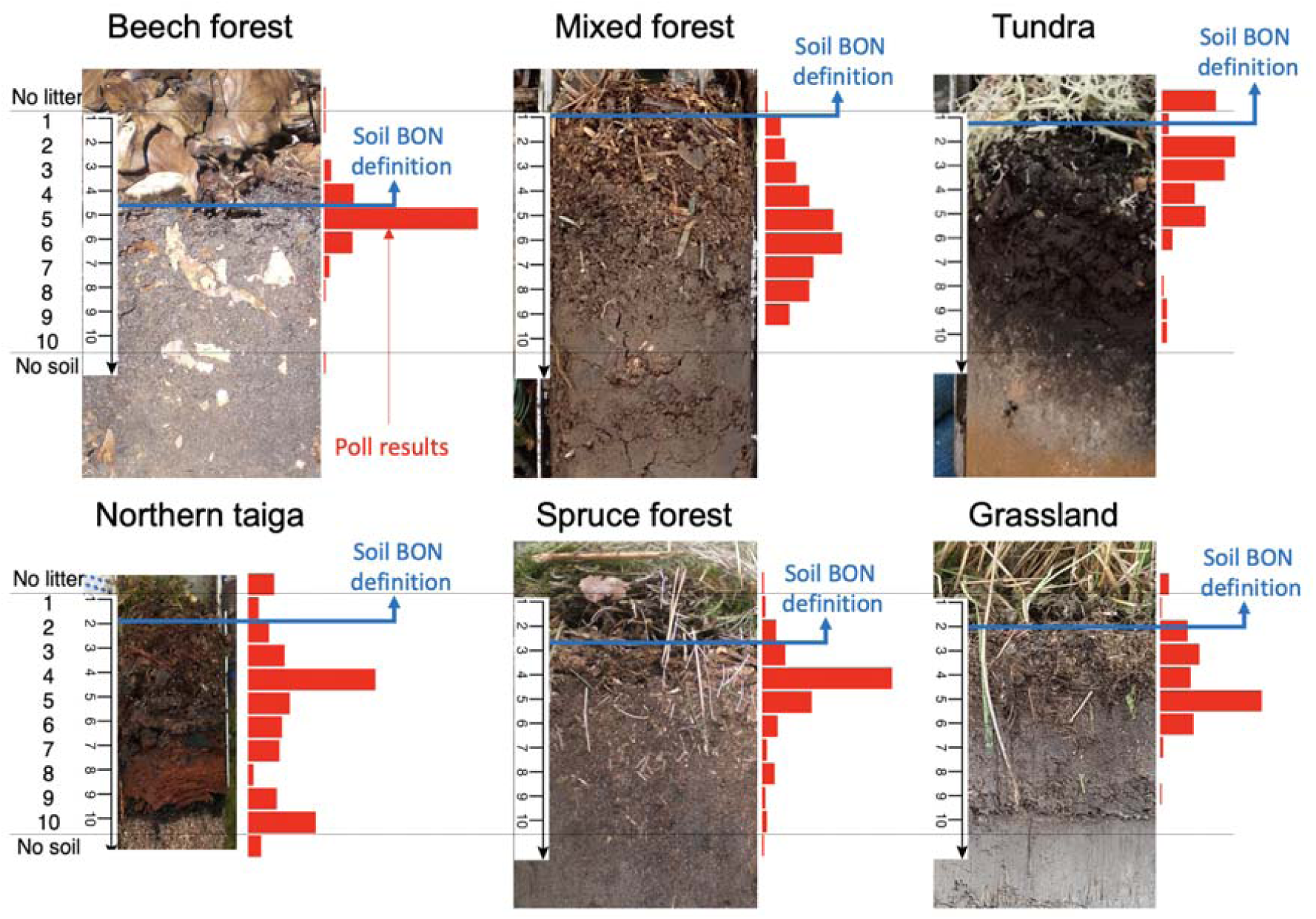
Variation in the definition of ‘litter’ by soil ecologists and the SBF Team solution. Results of a non-representative poll with the aim of finding the ‘litter-soil threshold’ on topsoil images are presented as red histograms. Responses were acquired anonymously from 170 soil ecologists after disseminating the poll through email contacts and Twitter, and represent an overview of opinions mostly from Europe (66% of respondents) and North and South America (27%). The poll was mostly completed by soil zoologists and functional ecologists (39% each). The SBF Team defines ‘litter’ as ***green vegetation remains, mosses, lichens and unfragmented dead leaves/wood with only little decomposition damage*** (100% organic matter, OL horizon, marked with blue lines and arrows) (Zanella, Ponge, Gobat, et al., 2018; Zanella, Ponge, Jabiol, et al., 2018).

#### Macrofauna collection

Large invertebrates (>3 mm in body length) including earthworms are picked up by hand from litter and soil, generally following the Tropical Soil Biology and Fertility method (TSBF) (Anderson, Ingram, 1993). We excavate soil monoliths 25 × 25 cm to a 10 cm depth and animals are hand sorted either in the field or brought to a laboratory. All social insects (ants, termites) regardless their size and all other animals belonging to the size class between 3 and 30 mm and not belonging to taxa of microarthropods (mites, pseudoscorpions, symphylans, diplurans, pauropods etc.) (Gongalsky, 2021) are collected and stored in 96% alcohol. Hand-sorting of macrofauna is preferred over dry extraction, because it has better compatibility with other existing global-scale initiatives (Lavelle et al. *under revision*), is conventionally used to collect earthworms, and catches less mobile animals like gastropods and insect larvae (Anderson & Ingram, 1993). Besides, large amounts of material do not have to be transported to the laboratory, which is hard for remote sites and may be disruptive.

### Animal extraction: *Local hubs*

Animal extractions are done at *Local hubs* to improve consistency of extractions across sites and make possible participation of research teams without extraction equipment. We distinguished *Wet extraction hubs* that are equipped with Baermann funnels and *Dry extraction hubs* that are equipped with dry Berlese-like extractors. These two types of hubs could (but do not have to) be at different locations and coordinated by different research teams. Each hub is expected to do extractions and animal imaging from 4-10 sites in the sampling year (i.e., 20-50 samples). *Local hubs* can receive central support from the SBF Team (e.g. imaging equipment and consultations regarding the extraction equipment and process), their coordinators are closely enrolled in the planning of sampling and add-on projects together with *National coordinators*, and have priority to lead regional-scale synthesis studies. The global assessment in the framework of the SBF Team can be combined with ongoing regional and local-scale studies as long as the sites are appropriate for both projects and the common sampling protocol is used. *Local hubs* are also responsible to organise the mid-term (at least 5 years) storage of samples, but are ***not*** expected to do field sampling (unless research team form both a *Local hub* and a *Field team*), animal sorting and identification (the latter is done from the images centrally at iDiv, Leipzig). A collaboration with museums for sample storage is encouraged.

#### Nematodes and enchytraeids extraction

These two groups are extracted separately using wet extractors. For both groups, we use Baermann funnels as the most commonly used and more accessible method in comparison to e.g. Oostenbrink elutriator (De Goede & Verschoor, 2000). Nematodes together with other microfauna are extracted through fine mesh filters to produce clean samples. We will follow a recent protocol for large-scale nematode assessments to improve extraction efficiency (Cesars et al., 2019). Formalin will be used as a storage agent to allow for potential future identifications.

#### Microarthropod extraction

To extract microarthropods, we use light bulb-equipped Tullgren/Berlese funnels (Karyanto et al., 2008; Tullgren, 1917) or Macfadyen/Kempson high-gradient extractors (Kempson et al., 1963; Macfadyen, 1961). These different extractor types provide a fair representation of community composition and biomass of microarthropods and, despite in some cases favouring different groups, yield comparable results (Andre et al., 2002; Edwards, 1991; Macfadyen, 1961). Since each laboratory is equipped with slightly different extractors, we will provide a protocol on the extraction procedure. The extraction is done through a 2 mm mesh to exclude large macrofauna, and ∼96% ethanol is used as the collection and storage solution to make the material suitable for potential genetic analyses.

### Animal identification

Soil communities will be characterised at each sampling site based on the list of taxonomic and functional groups with data on their abundance, body masses, and biomasses (the tentative list is given in Table 1). On the one hand, a taxonomic grouping should be detailed enough to make functional inferences as well as a food web reconstruction possible (Brussaard 1998; Briones 2014; Potapov 2021; Buchkowski & Lindo, 2021). On the other hand, the grouping should be generic enough to include all major taxa and regions and easy enough to allow the sorting by a general soil ecologist from a mixed community image, i.e. under a low magnification microscope. In most cases, we follow a taxonomic classification, because it allows for unambiguous grouping of animals and because functional roles and trophic niches of soil animals are in general related to their taxonomic position (Cardoso et al., 2011; A. M. Potapov et al., 2019). In several functionally diverse groups, such as mites, flies, ants, and termites broad taxonomic resolution may lead to information loss (Eggleton & Tayasu, 2001; Frouz, 1999; King, 2016; Schneider et al., 2004), but is dictated by the absence of appropriate taxonomic expertise across many regions. We are planning to approach these limitations in the follow-up molecular and taxonomic projects by establishing thematic collaborative networks. For instance, nematodes are planned to be sorted to trophic groups (Table 1) in collaboration with experts. In perspective, linking the global assessment of soil biodiversity with networks of taxonomic experts and organisation of taxonomic training programmes would allow for a great progress in the understanding of global soil biodiversity.

**Table 1.**
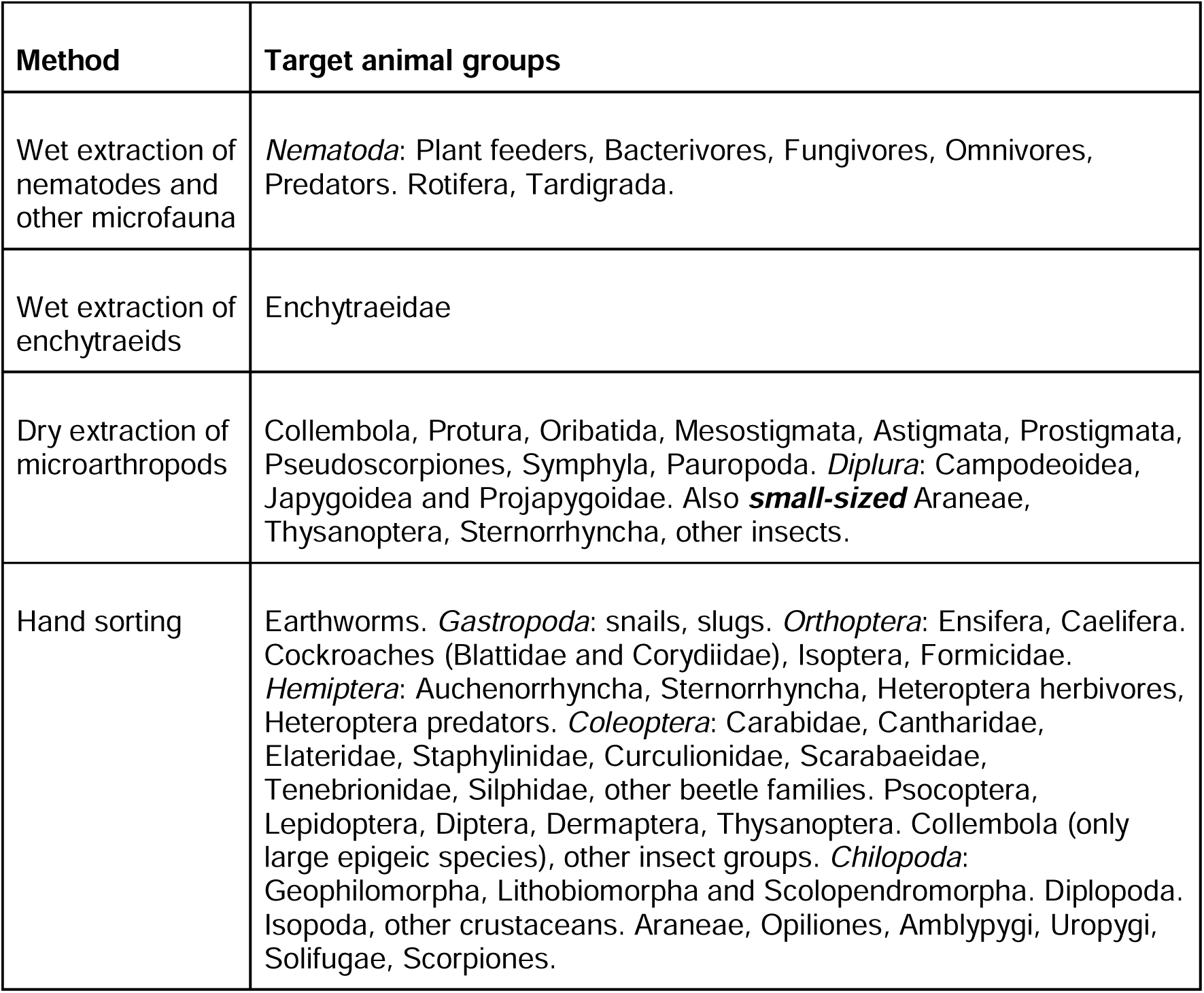
The list of animal groups which are counted and measured. Both taxonomic and functional groups are used for sorting. Individual groups are divided by comma. ‘Parent’ groups that include several target groups are given in *italics* and their ‘children’ are separated with full stop from other groups.

| Method | Target animal groups |
| --- | --- |
| Wet extraction of nematodes and other microfauna | *Nematoda*: Plant feeders, Bacterivores, Fungivores, Omnivores, Predators. Rotifera, Tardigrada. |
| Wet extraction of enchytraeids | Enchytraeidae |
| Dry extraction of microarthropods | Collembola, Protura, Oribatida, Mesostigmata, Astigmata, Prostigmata, Pseudoscorpiones, Symphyla, Pauropoda. *Diplura*: Campodeoidea, Japygoidea and Projapygoidea. Also **small-sized** Araneae, Thysanoptera, Sternorrhyncha, other insects. |
| Hand sorting | Earthworms. *Gastropoda*: snails, slugs. *Orthoptera*: Ensifera, Caelifera. Cockroaches (Blattidae and Corydiidae), Isoptera, Formicidae. *Hemiptera*: Auchenorrhyncha, Sternorrhyncha, Heteroptera herbivores, Heteroptera predators. *Coleoptera*: Carabidae, Cantharidae, Elateridae, Staphylinidae, Curculionidae, Scarabaeidae, Tenebrionidae, Silphidae, other beetle families. Psocoptera, Lepidoptera, Diptera, Dermaptera, Thysanoptera. Collembola (only large epigeic species), other insect groups. *Chilopoda*: Geophilomorpha, Lithobiomorpha and Scolopendromorpha. Diplopoda. Isopoda, other crustaceans. Araneae, Opiliones, Amblypygi, Uropygi, Solifugae, Scorpiones. |

### Animal identification and measurement

Animal sorting and identification is probably the most difficult and laborious part of soil animal assessments. Rapidly developing metabarcoding and metagenomic approaches are often seen as an alternative to visual identification (Liu et al., 2020; Oliverio et al., 2018); however, they have strong limitations in the light of our aims: (1) large-scale assessments using these approaches are expensive and require complex local facilities, thus are not inclusive for research teams/regions with limited resources; (2) mass-sequencing methods are non-quantitative due to extraction and amplification biases (Dopheide et al., 2019; Pereira-da-Conceicoa et al., 2021); (3) due to a limited trait data availability, it is not possible to extract body sizes for all soil invertebrate taxa based on genetic sequences and thus reliably calculate energy fluxes; (4) barcode libraries are absent for vast majority of soil invertebrate species to match phylogenetic positions with traits such as body size.

As an alternative high-throughput approach, we adopt image analyses of mixed community samples. Image analysis is based on visual animal identification and allows for direct body size estimations which can be used to estimate biomasses and energy fluxes. To acquire high resolution photographs, we will use a recently tested imaging pipeline for soil communities using a flatbed scanner (Potapov et al. *unpublished data*). This approach lowers the costs of imaging equipment to ca. 350 USD, thus being inclusive for teams with limited resources (Fig. 5). The imaging is done at *Local hubs* and all images are stored on central servers at iDiv, Leipzig.

**Fig. 5.**
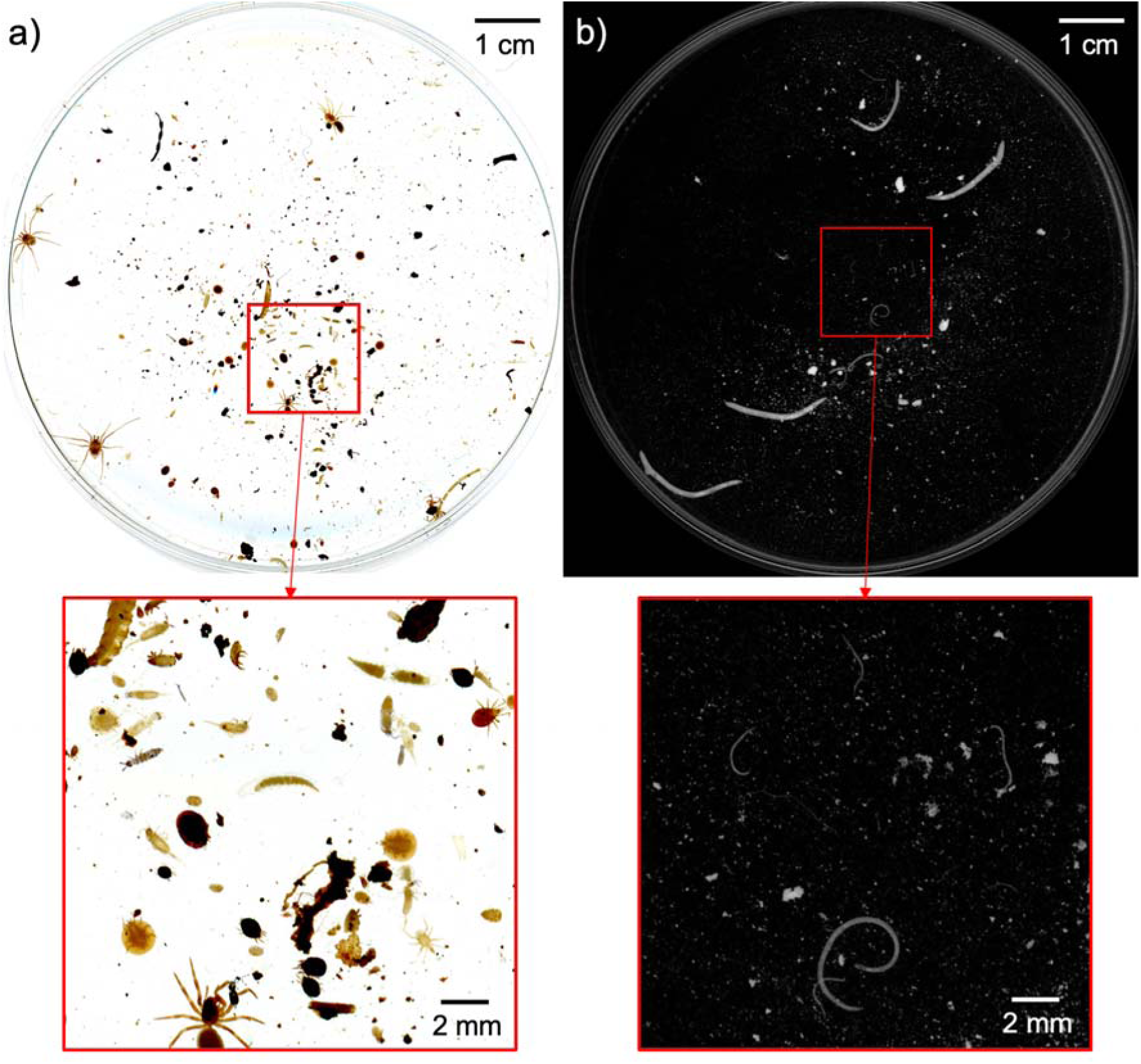
Images of mixed communities of soil arthropods (a) and nematodes (b) made with a flatbed scanner. The images have been taken from a Petri dish with 96% ethanol with a 4800 dpi resolution using Epson Perfection V600 (Seiko Epson Corporation, Japan). Close-up frames show that even small mites and nematodes can be detected on the images.

### Data acquisition

High-resolution pictures will be processed by the *Central team* using manual image annotations to develop a computer-vision pipeline based on deep learning algorithms (RCNN) (Sys et al., *under review*). Preliminary tests on a community of 10 springtail and two mite species showed a detection rate of 77% with a species identification accuracy of 90%. We expect improvement of the two metrics with further developments of the training library. Manual annotations will be used at the initial stages and then deep learning algorithms will be applied to streamline the identification and measurement processes. With this approach, all individuals of micro-, meso-, and macrofauna will be identified to the group level (Table 1) and measured. Individual body sizes will be converted to body masses and total community biomass using allometric equations (Newton & Proctor, 2013; Sohlström et al., 2018; Andrassy, 1956; Petersen, 1975). With development of a training image library, scanning and image analysis are expected to speed up group-level identification and body size estimations of soil invertebrates and considerably reduce workload of *Local hubs*. Our methodology can be applied by most soil ecology laboratories across the globe and will produce reliable and comparable data on density, biomass, and size distributions of all key functional groups of soil micro-, meso- and macrofauna found in the topsoil. All images and collected data will be centrally stored and accessible for the data providers who will have priority of analysing and publishing these data. Collected data and developed machine learning algorithms will be made openly available through public online platforms upon publication for the use of the research community.

## Future prospects

Data collected by the SBF Team will be linked to the data on soil, climatic, and microbial parameters, and relationships between soil animals and ecosystem functioning and their responses to environmental factors and human activities will be assessed. To analyse the animal data, we will use different approaches, such as path analysis (Eisenhauer et al., 2015), geospatial modelling (van den Hoogen et al., 2021), food-web reconstruction and modelling (A. Potapov, 2021), and energy flux approaches (Barnes et al., 2018; Jochum et al., 2021). We also plan to link collected animal data with multiple functional traits to assess global variation in the functional diversity of soil animal communities (Brousseau et al., 2018; Pey et al., 2014) and work on integration of our results in suggested animal-based biogeochemical models (Chertov et al., 2017; Deckmyn et al., 2020; Flores et al., 2021). Finally, any SBF Team participant can propose further ideas of how to use the collected data to the *Global coordination team* and test his/her/their hypotheses.

The developed protocol can be used beyond the SBF Team in any other compatible observational or experimental study that aims to accumulate standard and comprehensive data on soil animal communities across environmental and biotic gradients. To ensure the long-term data safety, compatibility, and accessibility, common databases on community images and animal counts will be established. We intend to upload collected data to Edaphobase (Burkhardt et al., 2014) which will make it publicly available directly and through the linked Global Biodiversity Information Facility (GBIF) (Heberling et al., 2021). To ensure the safety of collected materials, all animals will be stored in ∼96% ethanol under cool conditions (+4°C) for at least five years after the sampling, and this period is likely to be prolonged. These storage conditions will allow us to run potential add-on projects on biodiversity-related questions, such as metabarcoding of soil animal communities and taxonomic identification of selected animal groups. Established collaborative networks at the global and regional scales will serve as the coordination basis for such add-on projects, facilitating soil animal ecology and providing science-based evidence to policymakers to support soil biodiversity conservation and functioning of the terrestrial biosphere.

## Supporting information

Supplementary protocol

## Acknowledgements

AP and SS acknowledge support of the German Research Foundation (DFG–SFB 990, 192626868 – collaborative German - Indonesian research project CRC990 - EFForTS). NE acknowledges support of iDiv funded by the German Research Foundation (DFG– FZT 118, 202548816). GGB acknowledges support of CNPq (Process 310690/2017-0). XS acknowledges support of the National Natural Science Foundation of China (No. 42021005) and National Science & Technology Fundamental Resources Investigation Program of China (No. 2018FY100300). We thank all anonymous ‘Where is the litter?’ poll participants.

## Supplementary materials

Supplementary protocol is available for this paper.

