## Supplementary protocol for "Global monitoring of soil animal communities using a common methodology"

### Soil animal sampling - protocol overview

**THIS PROTOCOL IS TENTATIVE; CONTACT THE SBF TEAM**
**SOIL BON FOODWEB INFO**: https://soilbonfoodweb.org
**SOIL BON CORE INFO**: https://www.globalsoilbiodiversity.org/soilbon
This protocol has been developed by the Soil BON Foodweb Team to sample soil animals from Soil BON *sites* (it is not the main Soil BON protocol). Each *site* represents a habitat. For the Soil BON site selection, please contact *National coordinators*. This protocol can also be applied beyond Soil BON to produce comparable data across studies. On each *site*, **five *sampling points* are assessed**: one at the georeferenced center of the *site*, and four at *sampling points* 15 m in each of the four directions, N, S, W, E from the center. All samples at each *sampling point* are taken randomly within one square meter area. In total, the following materials and information are collected from each *site*: (**A**) *Site* and sampling event description; (**B**) 5 soil cores for wet extraction; (**C**) 5 soil cores for dry extraction; (**D**) 5 vials with hand-sorted large macrofauna; (**E**) 5 vials with hand-sorted earthworms; (**F**) 5 plastic bags with litter samples for weighing; (**G**) 5 photographs of topsoil profiles.
**Assessment of one *site* by two persons takes c. 3-4 hours**.

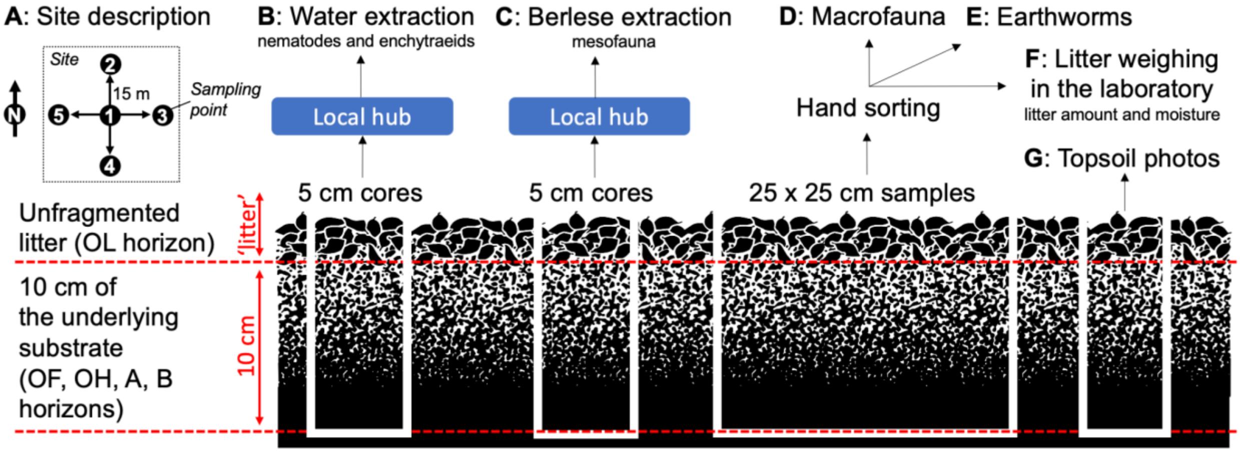



**Soil animal sampling - protocol overview** 1
- Materials for the field sampling [to print] 2
- Fieldwork overview cheatsheet [to print] 3
- A: Site and sampling event description [to print] 4
- B and C: Samples for water and Berlese extraction 5
- D, E and F: Hand-sorting of macrofauna and litter collection 6
- G: Topsoil photographs 7
- H: Pitfall traps (auxiliary) 8
- I and J: Deep soil macrofauna and earthworms (auxiliary) 8

**Wet extraction hubs (wet hubs)** **12**

**Dry extraction hubs (dry hubs)** **13**

**Imaging of animal communities** **14**

**Data acquisition** **15**

**KoBoToolbox for site data upload** **16**

### Materials for the field sampling [to print]

##### GENERAL FIELD EQUIPMENT

- Tape-measure, 15 m
- 5 cm diameter soil corer or a PVC/metal tube
- A sharp knife to cut soil
- A ruler to measure sampling depth
- Flat-shape spade to dig soil samples
- 25 by 25 cm frame to measure area
- Secateurs to cut ground vegetation
- Plastic tray(s) to sort macrofauna
- Tweezers and brushes to catch fauna
- A phone with camera/digital camera
- Background laminated template with a printed ruler for photographs
- A pencil or ethanol-resistant marker
- A hammer to hammer cores (optional)
- A saw to cut large roots (optional)
- Headtorch (optional)
- Gloves for field work (optional)

##### LABORATORY EQUIPMENT

- Scales to weigh the litter (min 0.1 g precision)

##### FIELD EQUIPMENT PER *SITE*

- 10 containers (~500-1000 ml) with lids or bags for transportation of soil cores
- 5 vials (15 ml+) with ~96% ethanol for macrofauna
- 5 vials (15 ml+) with ~96% ethanol for earthworms
- 2-3 backup vials for large animals with ~96% ethanol
- 96% ethanol (~150-200 ml)
- 5 plastic bags (~20-40 l) for litter
- 5 strong plastic bags (~40-60 l) for soil
- Labels* for vials and containers (6/*sampling point*, 30/*site*)
- Laminated label with the site code (if the site is not labeled by Soil BON).

*Labels should be robust, ethanol/waterproof and go inside and outside of vials. Use a laser printer, or a pencil. Minimum information: Project ID (e.g. SBF = SBF Team), PI Surname, Site ID, Sample type (B / C / D / E / F / G) and *sampling point* number (1-5), Date (YYYYMMDD). Example, plot code, sample type and sampling point are emphasised in bold:

| SBF_Potapov_**BF3_B2**_20220710 |
|---|

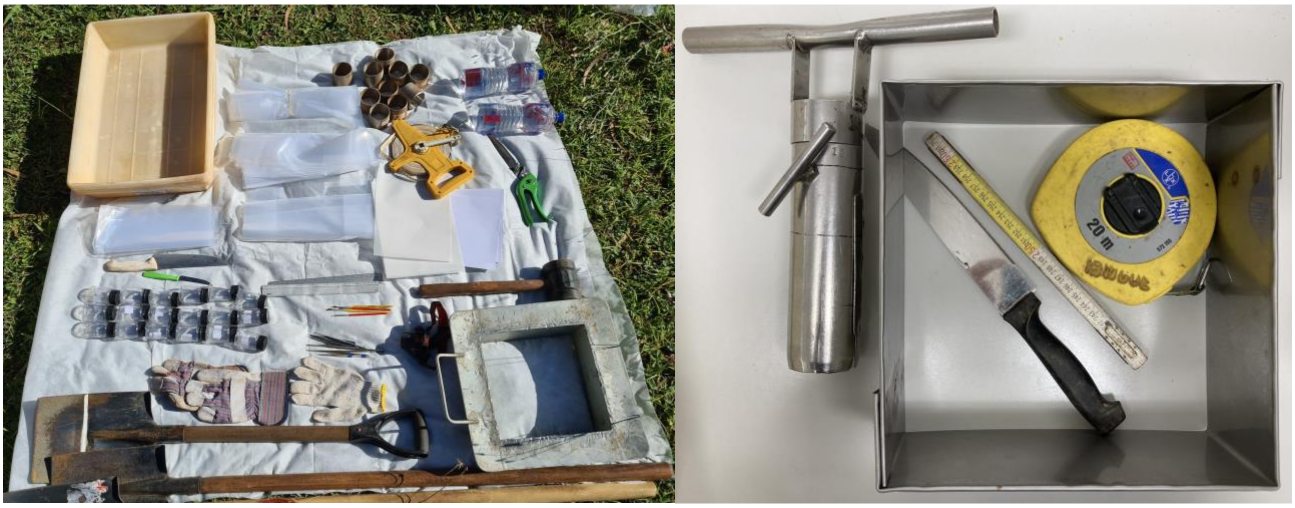

**Fig. 1** | Field sampling equipment. Example solutions, letters refer to items in the list.

### Fieldwork overview cheatsheet [to print]

**DO AS MUCH AS POSSIBLE OUTSIDE THE SITE TO NOT DISTURB IT.**

#### **A**: *Site* description

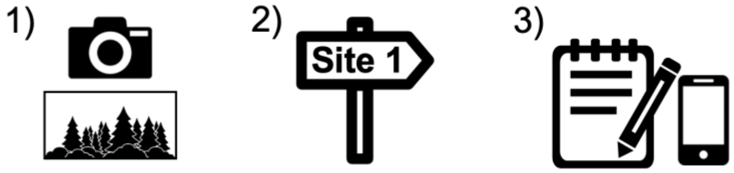


At each of the five sampling points:

#### **B and C**: Soil cores for wet and dry extractions

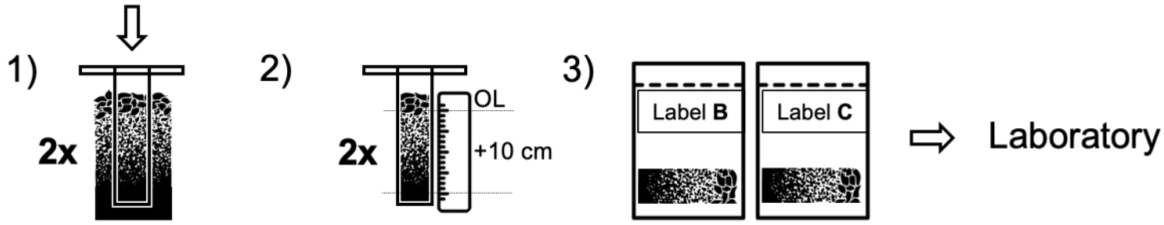


#### **D, E and F**: Hand sorting of large macrofauna and litter collection

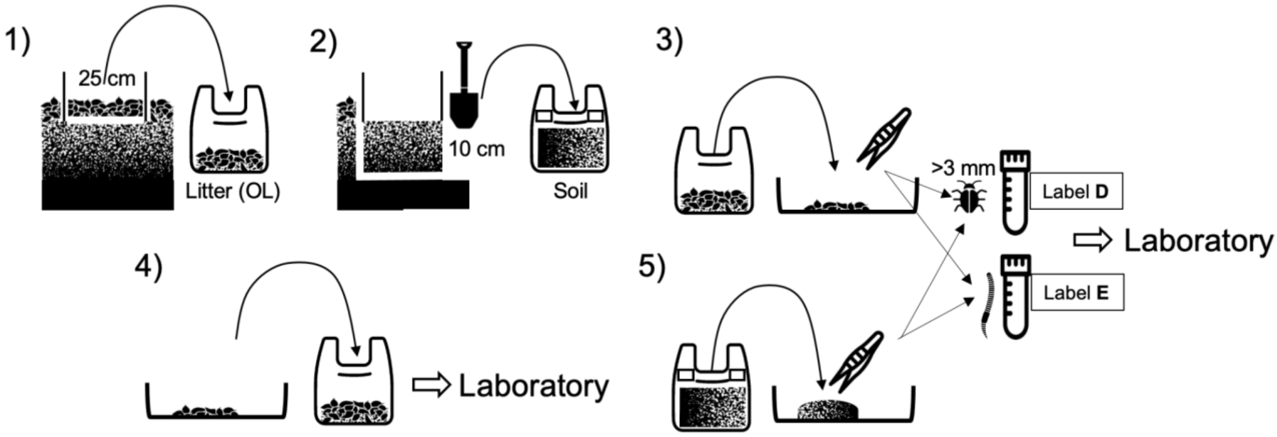


#### **G**: A photograph of topsoil profile

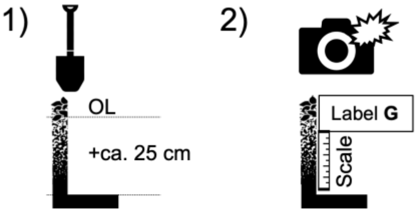

### **A**: Site and sampling event description [to print]

Make a photo of the entire sampling *site* overview with geolocalisation turned on (Fig. A1). Firmly fix the laminated label with the site code at the centre of the *site* (if it is allowed). Record the coordinates. Ensure to record complete *site*-level data in a table with exactly these columns, or upload this information via the submission system.

| | |
|---|---|
| ProjectID | ID of the project, fill “SBF” for the Soil BON Foodweb project |
| Researcher | Full name (first+second) of the PI, e.g. “Anton Potapov” |
| ContactEmail | Email of the person to contact regarding the data use |
| SiteID | ID of the *site*, should be unique to the project (avoid using “Site1” etc) |
| Country | Country name, e.g. “Nauru” |
| Locality | Nearest named geographical location, e.g. village or mountain name |
| DecimalLongitude | WGS84 coordinates in decimals, max 30 m uncertainty, e.g. 51.5377 |
| DecimalLatitude | WGS84 coordinates in decimals, max 30 m uncertainty, e.g. 9.9371 |
| SamplingDate | Date of the sampling in YYYYMMDD format, e.g. 20220710 |
| SamplingMethods | “ABCDEFG”=core methods, add “H”, “I”, “J” for auxiliary methods |
| Habitat | One from the list: “forest”, “grassland”, “shrub”, “agriculture”, ... |
| ProtectedArea | Fill “YES” for officially protected and “NO” for unprotected |
| CurrentVegetation | Free sentence to describe the age and type of the dominant vegetation |
| LanduseHistory | Known info on the site history in the last ~50 years; ecosystem age |
| Comments | Deviations from the protocol, if any. Any further relevant information |

**FIELD CHECKLIST**

- **A**: A photo of the *site* and *site* information
- **B**: 5 soil cores for water extraction
- **C**: 5 soil cores for Berlese extraction
- **D**: 5 vials with hand-sorted large macrofauna
- **E**: 5 vials with earthworms
- **F**: 5 plastic bags with litter samples for weightning
- **G**: 5 photographs of topsoil profiles

**POST-FIELD CHECKLIST**

- Cores+vials transported to *Local hubs* within 1 week
- *Site* and sampling event descriptions were submitted
- Photographs were named (see labels) and submitted
- Fresh and dry weights of litter were submitted

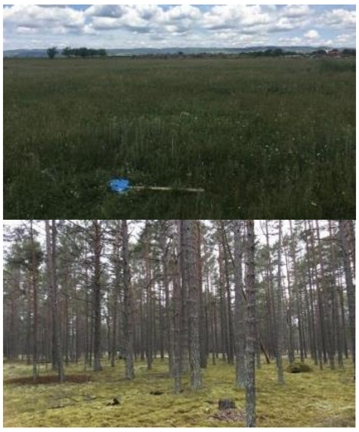

**Fig. A1** | Examples of the sampling *site* overview photographs

### **B** and **C**: Samples for wet and Berlese extraction

Target groups: nematodes, enchytraeids, microarthropods, small macrofauna (~10 mins/point)

Ten cores (1 for wet extraction and 1 for dry extraction at each sampling point) 5 cm in inner diameter, are taken with a corer/PVC/metal pipe or a 5 x 4 cm rectangle is cut (~20 $cm^2$ area). The sample includes ***litter*** (green vegetation remains, mosses, lichens and unfragmented dead leaves and wood = OL horizon, if present; Figs. 1 and B1) and the **underlying soil to a depth of 10 cm** from the *litter-soil interface* (OF+OH+A+B horizons). In shallow soil, cores are taken down to the maximum possible depth. If bulk cores are not feasible to sample (e.g. rocky soils), litter can be collected by hand from a 5 x 4 cm area and the underlying soil (excluding rocks) can be extracted with a knife to a depth of 10 cm to fill 200 ml volume.

1. Cut and **remove the ground vegetation** at the ground surface at the sampling spot; keep mosses and lichens.
2. **Take a soil core** down to a sufficient depth. If using a short corer/pipe, two or three layers are sampled sequentially and combined.
3. **Measure 10 cm** from the litter-soil interface (Fig. B1) in the sampling device. Remove and discard the deeper horizons.
4. Put the remaining *litter* and underlying soil in a **plastic container/bag**. Try to preserve the soil structure as much as possible.
5. Put a **label** inside, write the sample code outside (e.g. BF3_**B2**).
6. **Close the container/bag** and prepare it for transportation (a cool box is preferred).

All cores must be transported to the laboratories for extraction as soon as possible (one week maximum). Five cores are transported to the local **Wet extraction hubs** for wet extraction, five cores to the local **Dry extraction hubs** for dry extraction. Personal delivery is preferred; postal deliveries should be tested beforehand and materials packed appropriately. During the storage and transportation soil samples **should not be pressed, strongly shaken, overheated or overdried**; avoid direct sunlight.

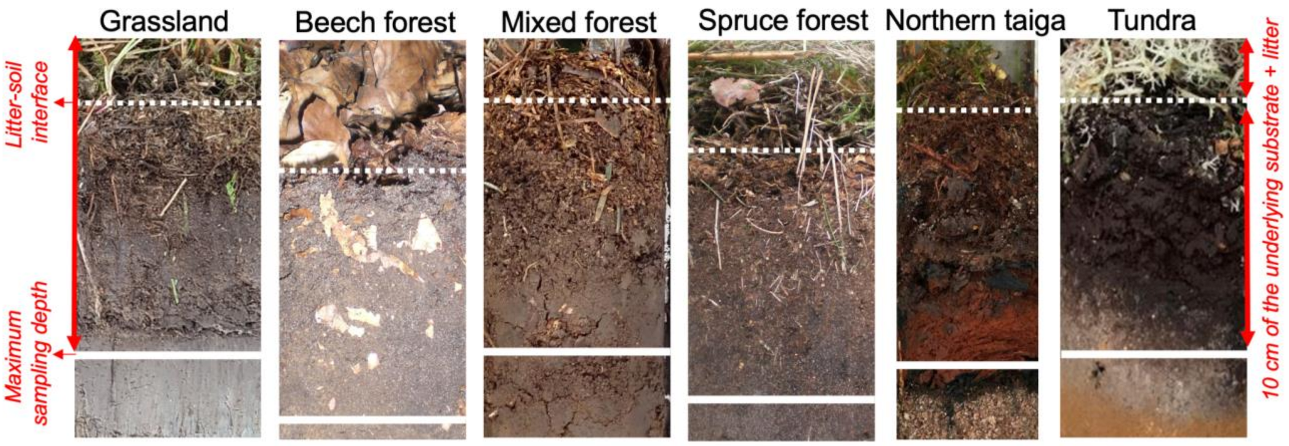


**Fig. B1 |** Examples of topsoil profiles with sampling depths marked. Each sample includes the entire litter layer + 10 cm of the underlying substrate below the *litter-soil interface* (i.e. 'soil'). Dotted white lines show *litter-soil interfaces*; solid white lines show maximum sampling depths.

### **D**, **E** and **F**: Hand-sorting of macrofauna and litter collection

Target groups: Large macrofauna, earthworms, social insects, litter weight (~60 mins/point/pers)

Five soil samples (1 at each sampling point) 25 x 25 cm (625 cm2), are taken with a spade. The sample includes ***litter*** and the **underlying soil to a depth of 10 cm** (see Fig. B1). All macroinvertebrates (≥ 3 mm in body length) are hand-sorted using tweezers or paint-brushes and put in a vial with ~96% ethanol. Earthworms are placed in a separate vial with ~96% ethanol. Mesofauna taxa except very large specimens (e.g. springtails ≥ 3 mm in body length) are ignored (Table 1). Hand sorting takes **a minimum time of 30 minutes per sampling point (two persons) and until the entire sample is checked**. Use a headtorch in limited light conditions. The animal collection must be done **outside of the *site*** to avoid disturbance of the *sampling points* and adjacent areas. It can be done in the laboratory/field station if litter and soil can be safely transported there within a day.

1. Put the **25 x 25 cm frame** on the ground and press it down to fix.
2. Cut and **remove the ground vegetation** within the frame; keep mosses+lichens.
3. **Collect *litter*** (+fauna) inside the frame with your hands and place it in a plastic bag. Wear gloves if dangerous animals are present in the area.
4. **Excavate the underlying soil** (10 cm) with a spade and put it in another bag.
5. **Sort fauna from the collected litter** by placing small amounts from the bag into a sorting tray. Collect macrofauna ≥ 3 mm in body length and earthworms in the vials.
6. **Place the litter back** in the plastic bag after all animals are captured. Close the bag, put the corresponding label inside and write the sample code on the bag.
7. **Sort the collected underlying soil** by placing small amounts from the bag into a sorting tray and breaking soil aggregates. Collect macrofauna ≥ 3 mm in body length and earthworms in separate vials. Animals from litter and soil are bulked together. Put the checked soil back and leave the site minimally disturbed.
8. Put a **label** inside both vials (E for earthworms and D for other macrofauna).
9. Transport and submit the vials with animals to the local **Dry extraction hub**.
10. Transport the litter in plastic bags (5 samples) to your laboratory.
11. **Back at the laboratory**:
    a. Weigh each litter sample with at least 0.1 g precision (=fresh weight).
    b. Air-dry all 5 litter samples separately in well-ventilated room until they are completely dry (>48 hours)
    c. Weigh each litter sample again (=dry weight).
    d. Submit both fresh and dry weight to the *Central team*.

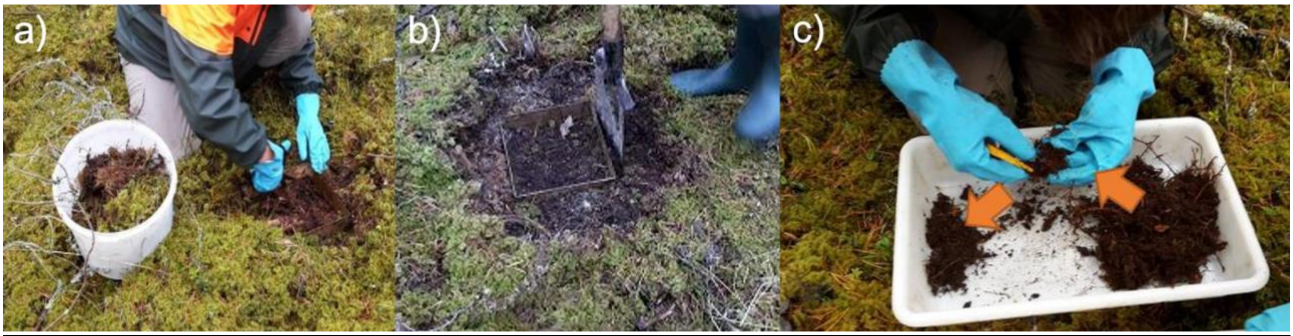


**Fig. D1 |** Hand-sorting of macrofauna: Litter removal within the 25 x 25 cm frame (a); Excavation of the soil monolith (b); Checking of substrates and animal collection (c).

### **G**: Topsoil photographs

Target variables: Diagnostic horizons, soil type and morphology (~5 mins/point)

5 photographs of topsoil profiles (1 at each sampling point) are taken using a phone camera **with a flashlight on** (Fig. G1). The photo is taken in the pit excavated for macrofauna collection and should include the full ***litter*** layer and **the underlying soil to a depth of 20 cm**, or down to the maximum possible depth if the soil is shallow). Please, turn on geolocalisation on your phone to keep tracking the sample location.

1. **Excavate soil** from the macrofauna pit (D,E,F samples) down to a depth of ~25 cm.
2. **Slice one of the walls** in the excavated pit to make it vertical and flat.
3. **Place a ruler** (cm scale) on the cleaned wall, zero is the *litter-soil interface*.
4. **Place the corresponding label** on the cleaned wall near the ruler.
5. **Make a photograph** with a flashlight on.
6. **Upload all photographs** via the submission system (see p. 11).

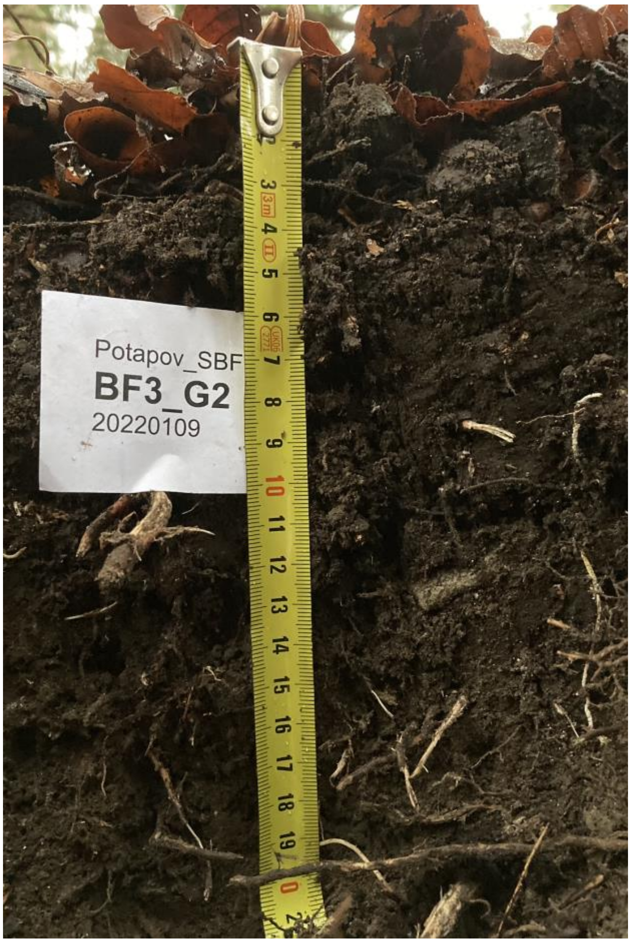


**Fig. G1 |** Example photograph of a topsoil profile with a template.

### **H**: Pitfall traps (auxiliary)

Target groups: Mobile macrofauna, social insects

Non-obligatory task, communicate with the *National coordinator* prior to implementation. Five pitfall traps (1 at each sampling point) are installed. Each trap is a jar with a diameter of 7.5 cm and a height of ca. 9 cm. Jars are dug into the soil so that the top edge of the glass is exactly level with the *ground surface* to avoid creating an obstacle for surface-active invertebrates. The traps are protected from rain and other disturbances by a roof (any waterproof material) and filled with 200 ml 75% propylene glycol. If there is high pressure from grazing livestock, the poles supporting the roof can be fitted with metal plates to prevent the roof from being trodden into the ground (Fig. H1). Pitfall traps are left in the field for 14 days. When collecting, the glass jars are closed with a lid and transported to the lab, where the invertebrates are rinsed with tap water and transferred to vials with ~96% ethanol for preservation. The five vials with collected fauna are submitted to the local **Dry hub** for imaging and storage.

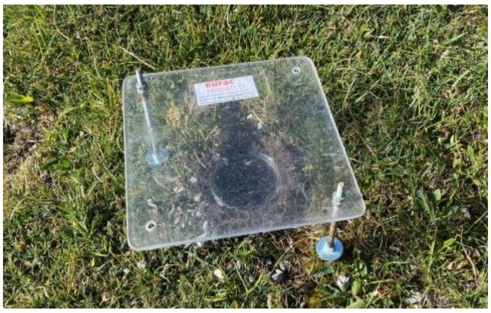

**Fig. H1 |** Installed Pitfall trap.

### **I** and **J**: Deep soil macrofauna and earthworms (auxiliary)

Target groups: Soil-living earthworms and other invertebrates

Non-obligatory task, communicate with the *National coordinator* prior to implementation. **J** and **I** samplings are incorporated in the core macrofauna sampling (**D**+**E**). After the top 10 cm of soil is removed, another underlying 10 cm are excavated and put in another plastic bag. The excavated soil is hand-sorted; macrofauna and earthworms are captured and preserved in ~96% ethanol in separate vials following the same approach as described above. This sampling results in two additional vials per *sampling point* and 10 additional vials per *site*. **J** and **I** samples are stored and processed separately from **D** and **E** samples. The 10 vials with collected fauna and earthworms are transported and submitted to the local **Dry hub** for imaging and storage.
